# Inactivation of NUPR1 promotes cell death by coupling ER-stress responses with necrosis

**DOI:** 10.1101/277384

**Authors:** Patricia Santofimia-Castaño, Wenjun Lan, Jennifer Bintz, Odile Gayet, Alice Carrier, Gwen Lomberk, José Luis Neira, Antonio González, Raul Urrutia, Philippe Soubeyran, Juan Iovanna

## Abstract

Genetic inhibition of NUPR1 induces tumor growth arrest. Inactivation of NUPR1 expression in pancreatic cancer cells results in lower ATP production, higher consumption of glucose with a significant switch from OXPHOS to glycolysis followed by necrotic cell death. Importantly, induction of necrosis is independent of the caspase activity. We demonstrated that NUPR1 inactivation triggers a massive release of Ca^2+^ from the endoplasmic reticulum (ER) to the cytosol and a strong increase in ROS production by mitochondria with a concomitant relocalization of mitochondria to the vicinity of the ER. In addition, transcriptomic analysis of NUPR1-deficient cells shows the induction of an ER stress which is associated to a decrease in the expression of some ER stress response-associated genes. Indeed, during ER stress induced by the treatment with thapsigargin, brefeldin A or tunicamycin, an increase in the mitochondrial malfunction with higher induction of necrosis was observed in NUPR1-defficent cells. Finally, activation of NUPR1 during acute pancreatitis protects acinar cells of necrosis in mice. Altogether, these data enable us to describe a model in which inactivation of NUPR1 in pancreatic cancer cells results in an ER stress that induces a mitochondrial malfunction, a deficient ATP production and, as consequence, the cell death by necrosis.

**Highlights:** NUPR1 expression promotes pancreatic cancer development and progression

NUPR1-depletion is a promising therapeutic strategy to be used for treating cancers

NUPR1-depletion induces ER stress, mitochondrial malfunction and a significant switch from OXPHOS to glycolysis followed by necrotic cell death

Inactivation of NUPR1 antagonizes cell growth by coupling a defective ER-stress response and a caspase-independent necrosis.

## Introduction

NUPR1 is a stress-inducible 82-aminoacids long, intrinsically disordered member of the AT-hook family of chromatin proteins. NUPR1 was first described as being activated in the exocrine pancreas in response to the cellular injury induced by pancreatitis (Mallo et al, 1997), an inflammatory disease, which in its chronic form, behaves as a preneoplastic condition for pancreatic cancer. Subsequently, the inducible expression of *Nupr1* was discovered to be a surrogate of the stress response caused by many stimuli in most cell types (Garcia-Montero et al, 2001) characterizing NUPR1 as a typical stress-associated chromatin protein. NUPR1 binds to DNA in a similar manner to other chromatin proteins (Encinar et al, 2001; Grasso et al, 2014) so as to control the expression of gene targets (Hamidi et al, 2012a). At the cellular level NUPR1 participates in many cancer-associated process including cell-cycle regulation, apoptosis (Malicet et al, 2006a; Malicet et al, 2006b), cell migration and invasion (Sandi et al, 2011), and DNA repair responses (Gironella et al, 2009). Indeed, NUPR1 has recently elicited significant attention due to its role in promoting cancer development and progression in the pancreas (Cano et al, 2014; Hamidi et al, 2012a). NUPR1-dependent effects also mediate resistance to anticancer drugs (Giroux et al, 2006; Palam et al, 2015; Tang et al, 2011), an important characteristic of this malignancy. We (Sandi et al, 2011; Vasseur et al, 2002b) and others (Emma et al, 2016; Guo et al, 2012; Kim et al, 2012; Li et al, 2017; Zeng et al, 2017) have shown that genetic inactivation of *Nupr1* antagonizes the growth of tumors in several tissues, including pancreatic cancer (Sandi et al, 2011) thereby supporting a role for this protein as a promising therapeutic target for the development of therapies for pancreatic cancer. Congruently, using a comprehensive approach that combines biophysical, biochemical, computational, and biological methods for repurposing FDA approved drugs in the treatment of pancreatic cancer, we have recently identified that the phenothiazine derivative, trifluoperazine, mimics the effect of the genetic inactivation of NUPR1, revealing its anticancer properties (Neira et al, 2017). The current study was designed to better understand the mechanisms by which targeting NUPR1 results in its tumor growth-inhibiting effects. We focused on determining the specific intracellular pathways that result in cell death after *Nupr1* inactivation *in vivo* (*Nupr1*-deficient mice) and *in vitro* (knockdown by either siRNA or CRISPR-Cas9). We found that in NUPR1-deficient cells, glucose consumption was switched from OXPHOS towards glycolysis resulting in a significantly reduced ATP production that promoted a caspase-independent programmed necrotic process. This defect was due to a mitochondrial malfunction, which in turn resulted from a strong ER stress. This report constitutes the first demonstration that inactivation of NUPR1 antagonizes cell growth by coupling two pathobiological cell phenomena, namely ER-stress response and caspase-independent necrosis.

## Results

### Genetic inactivation of NUPR1 induces pancreatic cell death by necrosis

In several *in vitro* and *in vivo* models of pancreatic cancer, NUPR1 inactivation inhibits the development and growth of this malignant tumor, highlighting the translational importance of this protein. However, the molecular mechanisms underlying these phenomena remain poorly understood. Previous work has demonstrated that *Nupr1* expression is rapidly and significantly induced by endoplasmic reticulum (ER) stress (Carracedo et al, 2006; Salazar et al, 2009). We therefore, evaluated the role of NUPR1 during ER stress by inhibiting its expression in ER-stressed cells. To carefully define this phenomenon, ER stress on pancreatic cancer cells (MiaPaCa2) was induced by using brefeldin A, thapsigargin or tunicamycin in combination with decreased the levels of NUPR1 using two different siRNAs (Figure S1). Subsequently, the necrotic and the apoptotic effects were measured through LDH release and caspase 3/7 activity, respectively. Upon ER stress induction, we found that LDH release was significantly higher in NUPR1 siRNA-transfected cells than in control cells (Figure 1A). Similarly, caspase 3/7 activity was also greater in NUPR1-depleted cells (Figure 1B). Combined, these experiments demonstrated that NUPR1 exerts both anti-apoptotic and anti-necrotic effects even in basal conditions, as well as during ER stress. Interestingly, pretreatment with the pan-caspase inhibitor, zVAD-FMK, did not prevent LDH release, as shown in Figure 1A, indicating that the role of NUPR1 to counteract necrosis is independent of its anti-apoptotic mechanism. We also used flow cytometry after co-labelling cells with Annexin V and PI to measure apoptosis and necrosis, respectively, in genetically NUPR1-depleted cells after brefeldin A, thapsigargin or tunicamycin treatment. Transfection with siRNAs against NUPR1 increased apoptotic and necrotic events, both in untreated and ER stressor-treated cells as presented in Figures 1C and D, which is in line with the data obtained concerning caspase 3/7 activity and LDH release. Inactivation of the *Nupr1* expression in Panc-1 cells (Figure S1), using a CRISPR-Cas9 knockout strategy, resulted in similar consequences, as shown in Figures 1E and F. Congruently with that data, after reconstitution of NUPR1 expression in CRISPR-Cas9 knockout Panc-1 cells by transfecting a NUPR1-expression vector, cell viability was almost completely rescued (Figures S2 and S3). Altogether, these data indicate that inactivation of NUPR1 increases the sensitivity of cells to apoptosis and to necrosis, and that the necrotic process is independent of caspase 3/7 activation. Thus, this data demonstrate that cell death after inactivation of NUPR1 occurs by a combination of both, apoptotic and necrotic processes.

**Figure 1.**
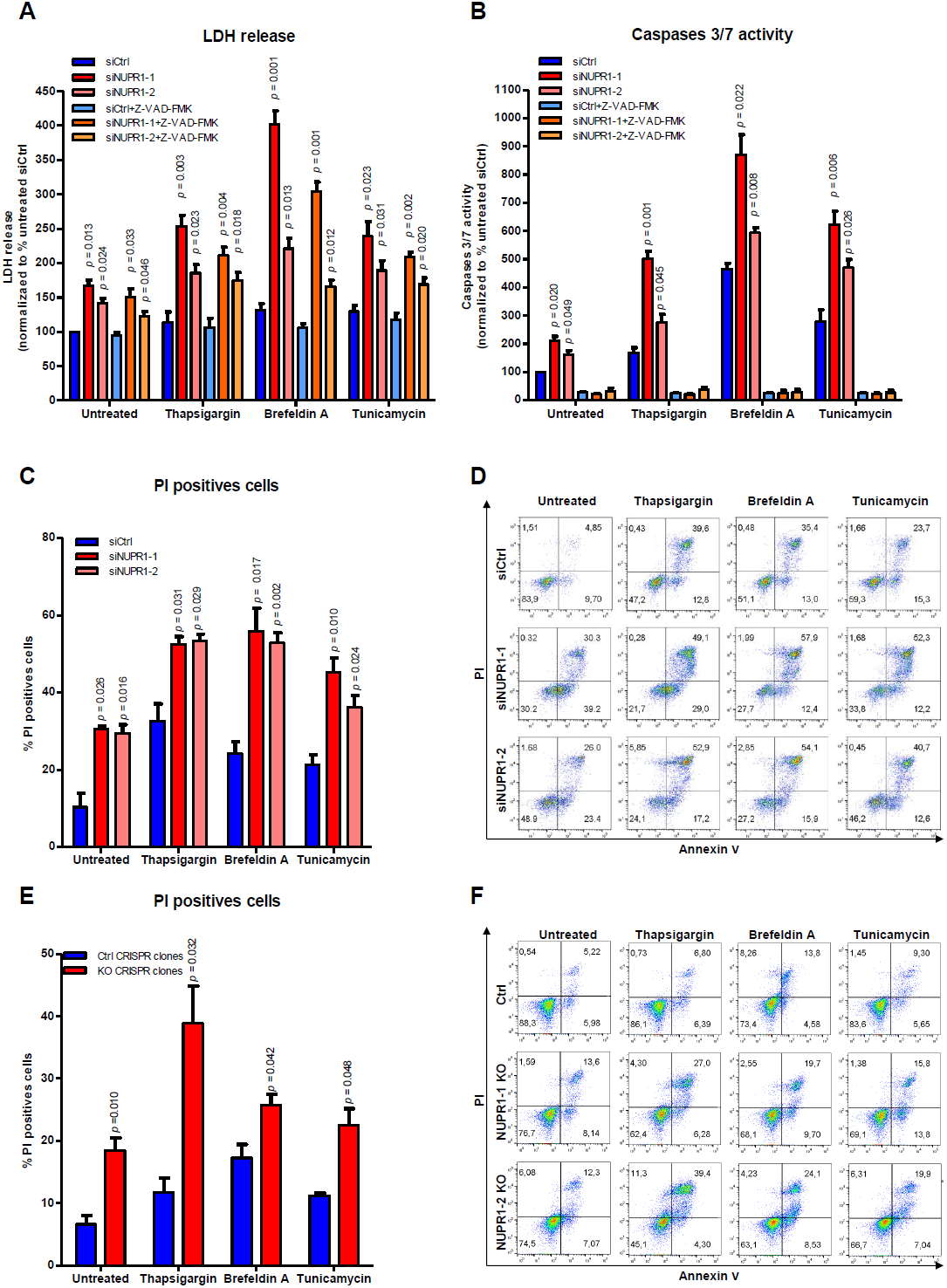
Nupr1 protects from cell death in stress situations *in vitro*. MiaPaCa2 cells were transfected with siCtrl or two different siNupr1 for 48 h. Cells were then incubated with thapsigargin, brefeldin A, or tunicamycin at 1 μM, for another 24 h in the presence or not of Z-VAD-FMK 10 μM. LDH release (A) and caspase 3/7 activity (B) were measured; data values were normalized to the siCtrl. In transfected MiaPaCa2 cells, flow cytometry analysis of annexin-V and PI staining following 24 h of treatment with thapsigargin, brefeldin A, or tunicamycin, showing % of PI-positive cells (C) and a representative experiment of the dot plot profile of cells (D). Data are means of triplicates ± SEM. For each treatment, statistically significant differences (only p values ≤ 0.05) from siCtrl are shown. Three Panc-1 control clones (with wild-type *NUPR1*) and 6 *Nupr1* knockout clones, developed using CRISPR-Cas9 technology, were cultured in DMEM and incubated with thapsigargin, brefeldin A, or tunicamycin at 1 μM for 24 h. Flow cytometry analysis was carried out after Annexin-V and PI staining. The percentage of PI-positive cells (E) and a representative experiment of the dot plot profile of cells (F) was showed. Data are means of triplicates ± SEM of 3 control and 6 *Nupr1* knockout clones. For each treatment, statistically significant differences (only p values ≤ 0.05) from control clones are shown.

### NUPR1 inactivation decreases ATP production

Since the normal production and handling of ATP as both an energetic and signaling-associated molecule is vital for cells (Golstein & Kroemer, 2007), we evaluated if ATP metabolism was implicated in the effects of NUPR1 inactivation on necrosis and apoptosis. Notably, we found that the levels of ATP were significantly reduced in cells transfected with *Nupr1* siRNAs, in untreated and in ER-stressed cells (Figure 2A). Again, inhibition of caspase 3/7 activity did not prevent this effect. Consequently, we considered that the decrease in the intracellular ATP content might be the result of reduced efficiency of the mitochondrial metabolism or decreased substrate availability for synthesizing these molecules within the cells (Kroemer & Pouyssegur, 2008). To address this issue, we therefore evaluated the mitochondrial oxygen consumption rate (OCR) at basal conditions as well as after treatment with an inhibitor of ATP synthase, oligomycin, or a potent mitochondrial oxidative phosphorylation uncoupler, carbonilcyanide p-triflouromethoxyphenylhydrazone (FCCP). Oligomycin treatment allowed the calculation of the OCR used by mitochondria exclusively for ATP production, while FCCP injection estimated the maximal respiration capacity of this organelle under stress situations. We measured the OCR in pancreatic cancer cells with *Nupr1* siRNA-mediated knockdown (Figure 2B) or inactivation by CRISPR-Cas9 (Figure 2C), as well as in pancreatic acini obtained from *Nupr1-KO* and *WT* mice (Figure 2D). Interestingly, compared with control cells, the OCR was strongly decreased in cells depleted of NUPR1 by siRNAs or CRISPR-Cas9-based knockout at basal respiration, as well as at the maximal respiration capacity (Figures 2B and 2C). In addition, we found that the OCR was drastically reduced in *Nupr1-KO* acini isolated from animals compared to those from WT mice (Figure 2D). Then, in order to evaluate if the oxidization of any of the main fuels used by the mitochondria for ATP production, which are glutamine, fatty acids, and glucose, could be compromised in NUPR1-inhibited cells, we used the Mito Fuel Flex Test. Measurements were obtained of the OCR for the basal respiration by the cells, and subsequently, we inhibited glutamine, fatty acid, or glucose metabolism for mitochondrial ATP production with BPTES, etomoxir, and UK5099 respectively. A decrease in the OCR after these methods of inhibition would indicate that the cells were metabolizing the substrate. Indeed, we found that after the injection of any of these three inhibitors, there was a decrease in the OCR (Figure 2E) in control cells. In contrast, in NUPR1-depleted cells, after the inhibition of the glutamine, fatty acid, or glucose oxidation, mitochondrial OCR was not modified as shown in Figure 2E. These results indicate that mitochondria are not appropriately oxidizing these substrates to generate ATP in the absence of NUPR1. Congruent with these results, we found that the expression of distinct enzymes involved in the tricarboxylic acid cycle, such as *SDHA, FH, PDHA*, and *IDH2*, was significantly downregulated in NUPR1-depleted cells as presented in Figure 2F. Since O_2_-dependent ATP production is inefficient in NUPR1-deficient cells, we evaluated the energy production by anaerobic glycolysis, measured as the extracellular acidification rate (ECAR) by the proton efflux into the media during lactate release. These experiments demonstrated that the ECAR increased under basal glycolysis and during inhibition of mitochondrial respiration after oligomycin treatment (glycolytic capacity) in NUPR1-depleted cells (Figure 3A), thereby resulting in lower cellular reserves of glucose. An increase in ECAR values was also observed with CRISPR-Cas9 knockout *Nupr1* cells, as shown in Figure 3B, corroborating this effect. Furthermore, in pancreatic acini obtained from *Nupr1-KO* and *WT* mice, we found that *Nupr1^−/−^* acini used more anaerobic glycolysis than *WT*, with a consequent reduction in the intracellular reserve of glucose (Figure 3C). This observation was also confirmed by directly measuring lactate release into the culture media by NUPR1-deficient cells (Figure 3D). We also detected a significant increase in glucose uptake in NUPR1-defficient cells (Figure 3E), whereas glutamine uptake remained unchanged (Figure 3F). Finally, measuring the OCR and the proton production rate by cells in the extracellular medium showed a rate of ATP production by glycolysis in control cells of 14.63 ± 1.30 pmoles/min/1000 cells, which represented 12.67% of the total ATP produced. For the siNUPR1-1-and siNUPR1-2-transfected cells, the rate was 22.75 ± 1.59 and 22.26 ± 0.80, respectively, which represented 32.43% and 30.74 % of the total ATP production (Figure 3G). Accompanying these events, we also found that several genes involved in glycolysis, such as *PDK, GPI, TPI, PGK1*, and *PGAM1*, increased their expression (Figure 3H). Taken together, these results strongly support a model whereby, under NUPR1 deficiency, the ATP production by glucose is switched towards glycolysis instead of OXPHOS. However, glycolysis is less efficient in terms of ATP production than OXPHOS, and therefore, the glucose reserve is rapidly consumed. Thus, NUPR1-depleted cells are limited by intracellular ATP availability, which contributes to the triggering of cell death by necrosis.

**Figure 2.**
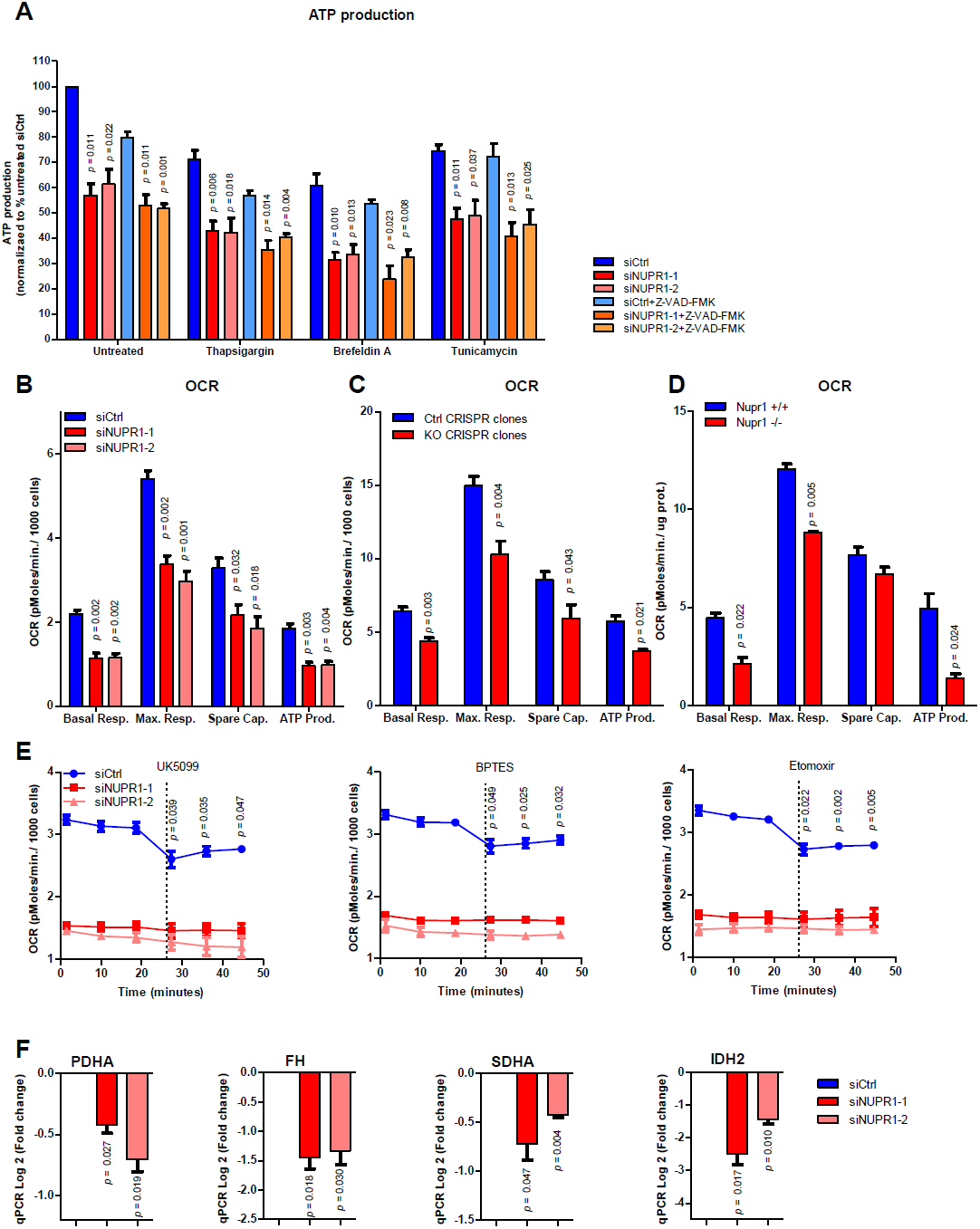
Nupr1 deficiency induces a decrease in ATP production by OXPHOS suppression. MiaPaCa2 cells were transfected with siCtrl or two different siRNA against *Nupr1* for 48 h. Cells were then incubated with thapsigargin, brefeldin A, or tunicamycin at 1 μM for another 24 h period in the presence or not of Z-VAD-FMK 10 μM, and ATP production (A) was measured; data values were normalized to the siCtrl. Mitochondrial respiration reflected by oxygen consumption rate (OCR) levels was measured in MiaPaCa2 cells transfected with siCtrl, siNUPR1-1 or siNUPR1-2 for 72 h (B). Mitochondrial respiration was also measured in three clones of Panc-1 control cells (with wild-type *NUPR1*), and six clones of *Nupr1* knockout cells, developed by CRISPR-Cas9 technology. Data are means of triplicates ± SEM of the 3 control and 6 knockout clones (C). Finally, we measured mitochondrial respiration in pancreatic acinar cells from WT (*Nupr1*^+/+^) and *Nupr1*^−/−^, mice (D). Only statistically significant differences (p values ≤ 0.05) were shown. The OCR was measured under basal conditions or following the addition of oligomycin, FCCP, or rotenone/antimycin A. The rates of OCR for basal respiration, maximal respiration, spare capacity, and ATP production were quantified as described in the Materials and Methods section. The OCR was again determined in MiaPaCa2 cells transfected with siCtrl, siNUPR1-1, or siNUPR1-2 when cells were challenged with UK5099, BPTES, or etomoxir; inhibitors of glucose oxidation, glutaminase, and carnitine palmitoyl-transferase 1A, respectively (E). Total RNA was extracted to monitor the mRNA levels of the genes involved in the Krebs cycle using RT-qPCR (F). Data are means of triplicates ± SEM. For each treatment, statistically significant differences (p values ≤ 0.05) from siCtrl are shown.

**Figure 3.**
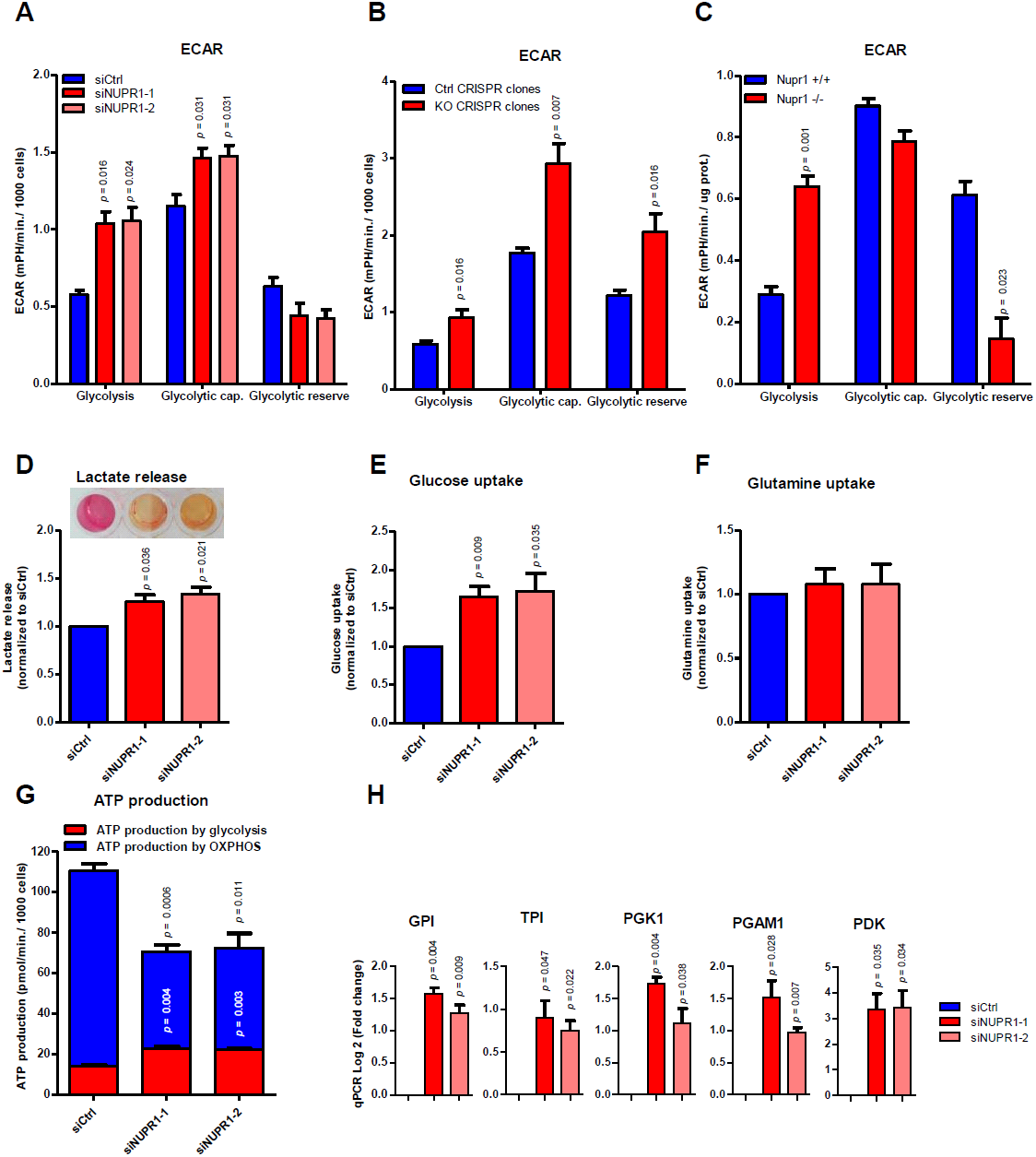
*Nupr1* deficiency promotes anaerobic glycolytic metabolism. Extracellular acidification rate (ECAR) levels were measured in MiaPaCa2 cells transfected with siCtrl, siNUPR1-1, or siNUPR1-2 for 72 h (A). The ECAR was measured in three clones of Panc-1 control cells (with wild-type *Nupr1*) and six clones of Nupr1 knockout cells, developed by CRISPR-Cas9 technology. Data are means of triplicates ± SEM of three control and six knockout clones (B). Statistically significant differences from Panc-1 control cells with p values ≤ 0.05 are shown. Also, the ECAR was measured in pancreatic acinar cells from WT (*Nupr1*^+/+^) and *Nupr1^−/−^* mice (C). The ECAR was measured under basal conditions or following the addition of glucose, oligomycin, or 2-Deoxyglucose. The rates of ECAR for glycolysis, glycolytic capacity, and glycolytic reserve were quantified as described in the Materials and Methods section. Lactate release (D), glucose (E) and glutamine (F) uptake and were measured in the extracellular medium after 24 h in culture. A representative image of DMEM collected after 72 h of culture is shown (D). ATP production by OXPHOS and anaerobic glycolysis were determined by the OCR and proton production rate, respectively (G); data values were normalized to the siCtrl. Total RNA was extracted to monitor mRNA levels of the genes involved in glycolysis using RT-qPCR (G). Data are means of triplicates ± SEM. For each treatment, statistically significant (p values ≤ 0.05) differences from *Nupr1*^+/+^ mice or siCtrl are shown.

### NUPR1 inactivation alters the function and localization of mitochondria

The results described above demonstrate that inactivation of NUPR1 induces a defect in mitochondrial OXPHOS activity, which is evidenced by the strong decrease in O_2_ consumption and ATP production, even if the cell attempts to compensate by activating the production of ATP through glycolysis. Consequently, we further evaluated the normal functioning of the mitochondria in cells in which NUPR1 had been genetically inactivated. First, we evaluated the proton gradient by measuring the membrane potential using the Mito ID system. In mitochondria, proton gradients generate a chemiosmotic potential known as the proton motive force which is used for the synthesis of ATP by OXPHOS, Ca^2+^ uptake, and antioxidant defense. The measurement of the chemiosmotic potential is a key indicator of mitochondrial and cell health or injury. In NUPR1-depleted cells, we found a significant depolarization of the mitochondrial membrane which is reflected by a decrease in the fluorescence of dye, as presented in Figure 4A and B, which explains, at least in part, the observed defect in ATP production. We then measured superoxide anion production by the mitochondria using the MitoSOX reagents, by both fluorescent microscopy and flow cytometry. These studies are important since superoxide anion generation by mitochondria triggers oxidative damage and contributes to retrograde redox signaling from the organelle to the rest of the cell. In this regard, both flow cytometry and fluorescence microscopy showed a significant increase in the intracellular levels of superoxide anions in NUPR1-deficient cells (Figures 4C and 4D). Thus, these experiments clearly demonstrate that NUPR1 inactivation affects the function of mitochondria, which explains the impact on ATP metabolism described above. The cellular distribution of mitochondria was analyzed in NUPR1-deficient cells using the MitoTracker Deep Red reagent counterstained with Alexa Fluor 488 phalloidin for cytoplasm and DAPI staining for the nucleus. The images shown that the distribution of these organelles remained localized in a limited perinuclear region, which differed from control cells, where mitochondria were more homogeneously localized throughout the cytoplasm as a structured network (Figure 5A). Transmission electron microscopy analysis of the mitochondria confirmed their limited perinuclear localization and the absence of significant ultrastructural changes of the organelle (Figure 5B). Through these experiments, we found no apparent change in the overall number of mitochondria in NUPR1-deficient cells (Figure 5C). However, these techniques revealed that mitochondria of the NUPR1-deficient cells were almost always associated with the ER, which was enlarged, as often observed during ER stress (Figure 5D). We therefore sought to gain insight into whether this observation indeed indicated that Nupr1-defficient cells were undergoing ER stress by measuring intracellular Ca^2+^ concentration using flow cytometry. Indeed, one of the most typical features of ER stress is the massive Ca^2+^ release from the stressed ER to the cytoplasm, which causes Ca^2+^ to move from the ER to mitochondria. We found a considerable increase in this parameter with values that resemble those found in our positive controls of ER Ca^2+^ release, namely brefeldin A-or tunicamycin-treated cells (Figure 5E). Concomitantly, we found a significant downregulation in the expression of two genes: *TFAM*, which encodes a key mitochondrial transcription factor that functions in mitochondrial DNA replication and repair, and *NRF1*, which encodes a transcription factor that activates expression of nuclear genes required for respiration and mitochondrial DNA transcription and replication (Figure 5F). In contrast, expression of *LONP1*, a gene encoding a mitochondrial protease involved in the selective degradation of misfolded, unassembled, or oxidatively damaged polypeptides in the mitochondrial matrix, and *PINK1*, a gene encoding for a serine/threonine protein kinase that is thought to protect cells from stress-induced mitochondrial dysfunction, were overexpressed (Figure 5F). Altogether, these results reveal a malfunction and abnormal subcellular distribution of mitochondria which may be due to the massive Ca^2+^ release from the stressed ER in NUPR1-deficient cells.

**Figure 4.**
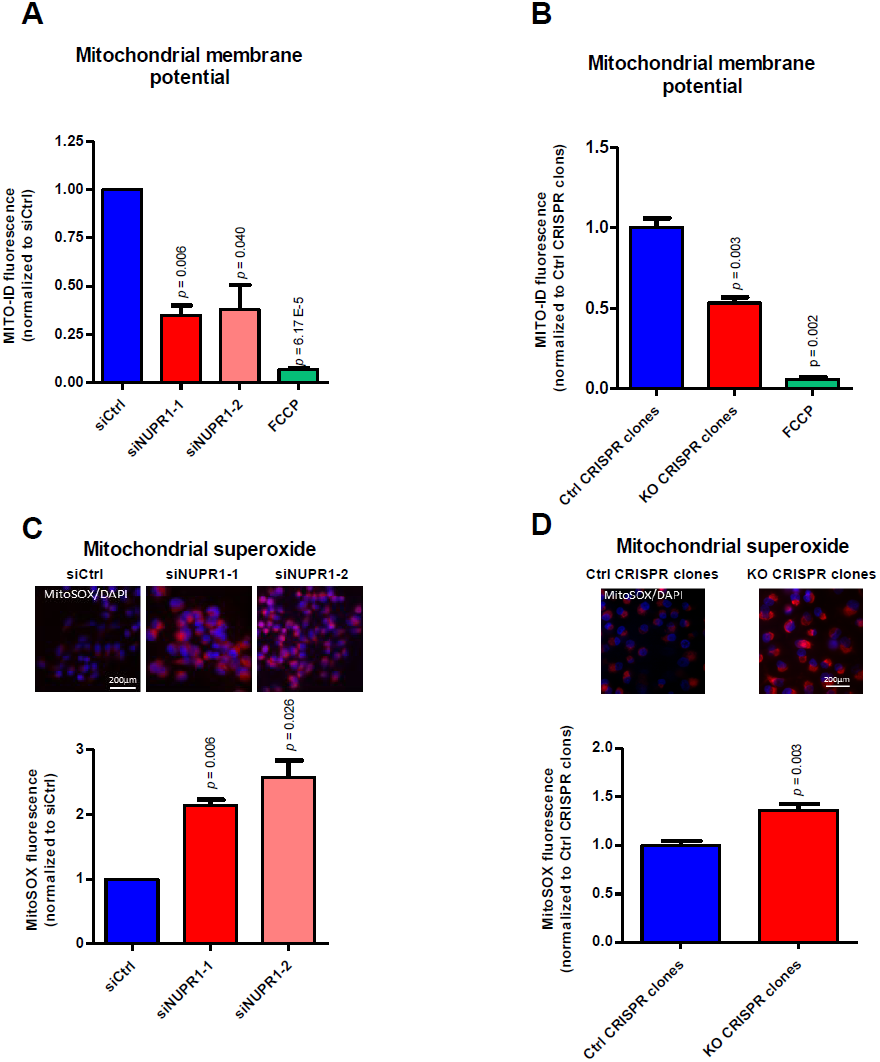
*Nupr1* deficiency leads to a mitochondrial membrane potential disruption inducing ROS overproduction. Flow cytometry analysis of MiaPaCa2 cells transfected with siCtrl, siNUPR1-1, or siNUPR1-2 for 72 h was carried out using MITO-ID for analysis of the mitochondrial membrane potential (A). MITO-ID was also applied to three clones of Panc-1 control cells (with wild-type *NUPR1*) and six clones of *Nupr1* knockout cells, developed by CRISPR-Cas9 technology. Data are means of triplicates ± SEM of the 3 control and 6 knockout clones (B). ROS production was detected using MitoSOX Red by flow cytometry analysis in the experimental cell groups as described above (C and D). Representative images of the fluorescence microscopy of both ROS indicators are shown (C and D). Data are means of triplicates ± SEM. Statistically significant differences from siCtrl with a p value ≤ 0.05 are shown.

**Figure 5.**
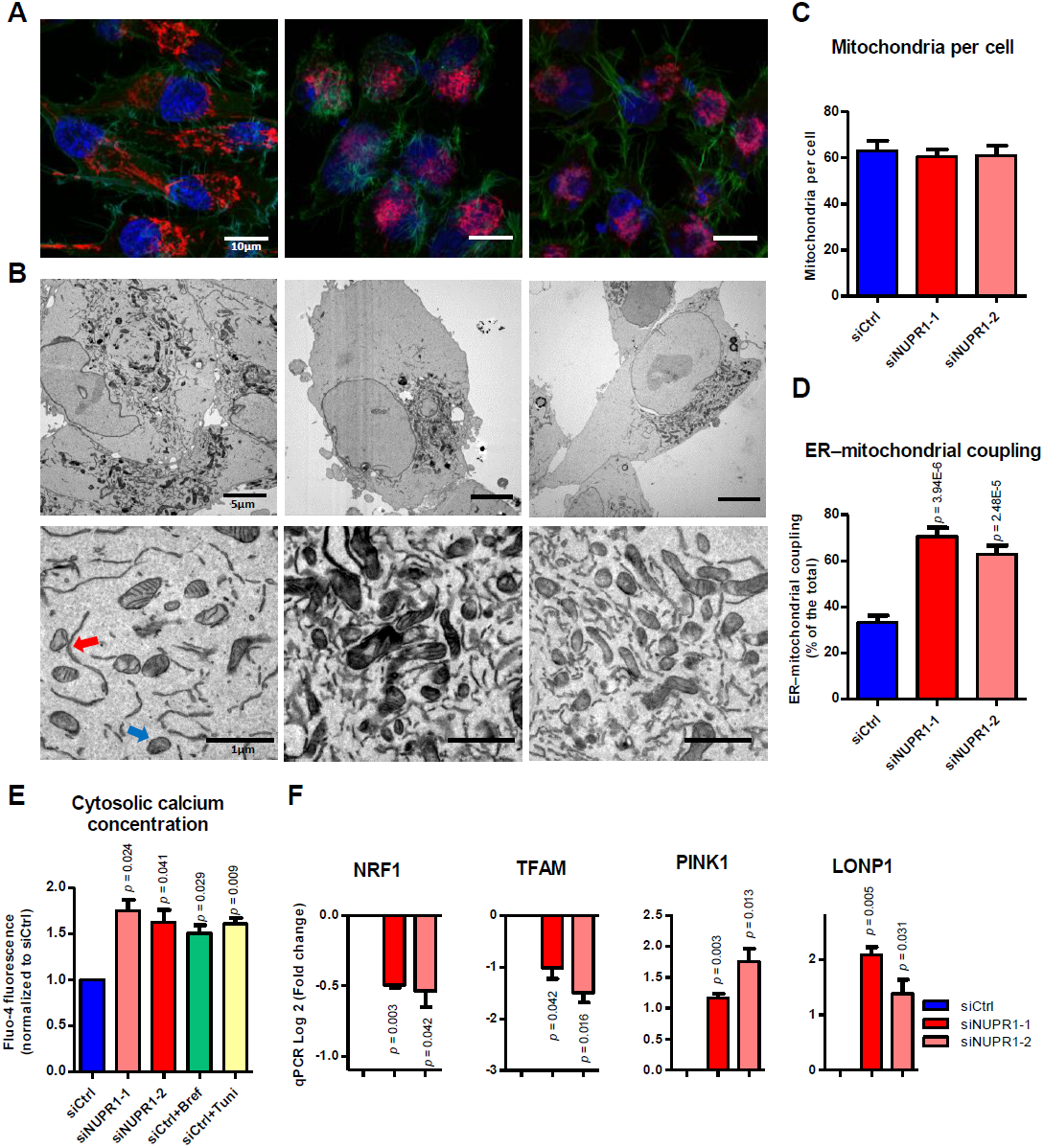
Malfunction of mitochondria in NUPR1-deficients cells. MiaPaCa2 cells transfected with siCtrl, siNUPR1-1 or siNUPR1-2 for 72 h were loaded with the mitochondrial selective probe MitoTracker Deep Red FM and, after fixation, marked with Alexa Fluor 488 Phalloidin and DAPI (A). Representative transmission electron microscopy images with different magnifications of the cells are shown. ER-mitochondrial contact or not are indicated by red or blue arrows, respectively (B). Mitochondria number per cell was evaluated counting them in transmission electron microscopy images; not less than 12 cells were counted for each condition (C). The percentage of mitochondria with ER contacts per field (mean of 10 fields) for each condition is shown; not less than 250 mitochondria were counted (D). Flow cytometry analysis was carried out using Fluo-4-AM for analysis of the cytosolic calcium concentration (E). Total RNA was extracted to monitor mRNA levels of the genes involved in the mitochondrial dynamic (F). Data are means of triplicates ± SEM. Statistically significant differences from siCtrl with a p value ≤ 0.05 are shown.

### NUPR1 inactivation induces failures in ER stress response

*Nupr1* expression is strongly induced in response to ER stress, but its role in ER stress response, if any, remains to be established. In fact, the ER stress response is accompanied by the activation of several ER stress-associated genes (Kato & Nishitoh, 2015), with unknown roles in this process for some of them. We therefore evaluated the role of NUPR1 in the ER stress response using several complementary approaches. The transcriptome of the MiaPaCa2 cells treated with a siRNA against *Nupr1* revealed a downregulation of several genes involved in ER stress-associated pathways when analyzed by both the Ingenuity Pathways Analysis (http://www.ingenuity.com/) and STRING (Szklarczyk et al, 2015) bioinformatics tools. The most significant biological pathways identified as inefficiently activated in NUPR1-deficient cells were: the response to unfolded protein (30 mRNA downregulated, FDR 7.53e-40), protein folding (28 mRNA downregulated, FDR=1.1e-32), the ER unfolded protein response (17 mRNA downregulated, FDR=1.55e-19), regulation of translational initiation (13 mRNA downregulated, FDR=2.95e-15), response to ER stress (16 mRNA downregulated, FDR 9.98e-14), the ER-nucleus signaling pathway (8 mRNA downregulated, FDR=1.48e-10), the ATF6-mediated unfolded protein response (6 mRNA downregulated, FDR=1.48e-10), the PERK-mediated unfolded protein response (6 mRNA downregulated, 1.33e-09), chaperone-mediated protein folding (8 mRNA downregulated, FDR=2.17e-09), response to stress (32 mRNA downregulated, FDR=5.97e-06), and the IRE1-mediated unfolded protein response (6 mRNA downregulated, FDR=2.14e-05) (see Table 1 and Table S1). These experiments, designed to directly evaluate the effects of NUPR1 in exocrine pancreatic cells, demonstrate that gene expression networks associated to ER stress are among the top pathways associated with expression changes under these conditions. To confirm these experiments *in vivo*, we made use of the *Nupr1-KO* mice model. Interestingly, transcriptomic analysis of pancreata from these mice analyzed using the same bioinformatics tools, also revealed an inefficiency in the ER stress response, for some of these biological processes, such as: the response to ER stress (10 mRNA downregulated, FDR=7.73e-12), response to unfolded protein (7 mRNA downregulated, FDR 6.82e-09), protein folding (7 mRNA downregulated, FDR=6.81e-07), ER unfolded protein response (5 mRNA downregulated, FDR=1.94e-06), cellular response to unfolded protein (5 mRNA downregulated, FDR=2.15e-06), and response to stress (12 mRNA downregulated, FDR=0.00255) (see Table 1 and Table S2). Thus, these results from our animal model, combined with those obtained in cells, show that NUPR1 deficiency mounts an inefficient ER stress response.

**Table 1.** Nupr1-deficiency induces a downregulation of the ER stress-associated pathways *in vitro* and *in vivo*.

| MiaPaCa2 treated with siNUPR1 |  |  |
| --- | --- | --- |
| Biological process | Downregulated genes | FDR |
| Response to unfolded protein | 30 | 7.53e-40 |
| Protein folding | 28 | 1.1e-32 |
| Endoplasmic reticulum unfolded protein response | 17 | 1.55e-19 |
| Regulation of translational initiation | 13 | 2.95e-15 |
| Response to endoplasmic reticulum stress | 16 | 9.98e-14 |
| Er-nucleus signaling pathway | 8 | 1.48e-10 |
| Atf6-mediated unfolded protein response | 6 | 1.48e-10 |
| Perk-mediated unfolded protein response | 6 | 1.33e-09 |
| Chaperone-mediated protein folding | 8 | 2.17e-09 |
| Response to stress | 32 | 5.97e-06 |
| Ire1-mediated unfolded protein response | 6 | 2.14e-05 |
| NUPR1<sup>-/-</sup> mice |  |  |
| Biological process | Downregulated genes | FDR |
| Response to endoplasmic reticulum stress | 10 | 7.73e-12 |
| Response to unfolded protein | 7 | 6.82e-09 |
| Protein folding | 7 | 6.81e-07 |
| Endoplasmic reticulum unfolded protein response | 5 | 1.94e-06 |
| Cellular response to unfolded protein | 5 | 2.15e-06 |
| Response to stress | 12 | 0.00255 |

Subsequently, we measured the expression of several genes involved in controlling the ER stress response, in NUPR1-depleted MiaPaCa2 cells, such as ATF6, IRE1α, PERK, EIF2A, XBP1, ATF4, CANX, CALR, BIP and HSP90 by RT-qPCR. Notably, some of these genes demonstrated significantly decreased expression in the absence of NUPR1 expression although not BIP and HSP90 (Figure 6A). In addition, expression of ATF6, IRE1α, PERK and BIP proteins was also measured by western blot on the siNUPR1-1-and siNUPR1-2-treated MiaPaCa2 cells. Interestingly, whereas levels of ATF6, IRE1α and PERK were decreased in NUPR1-depleted cells, BIP protein levels remained unaffected in strong correlation with its mRNA (Figure 6B). Altogether, these data suggest that expression of NUPR1 is necessary for an optimal ER stress response, which occurs, at least in part, through control of these genes at the transcriptional level. Therefore, when NUPR1 is inactivated, the ER stress response is suboptimal, with dramatic consequences on mitochondrial function and a strong deficit in ATP production, and as main consequence, cell death by necrosis is induced.

**Figure 6.**
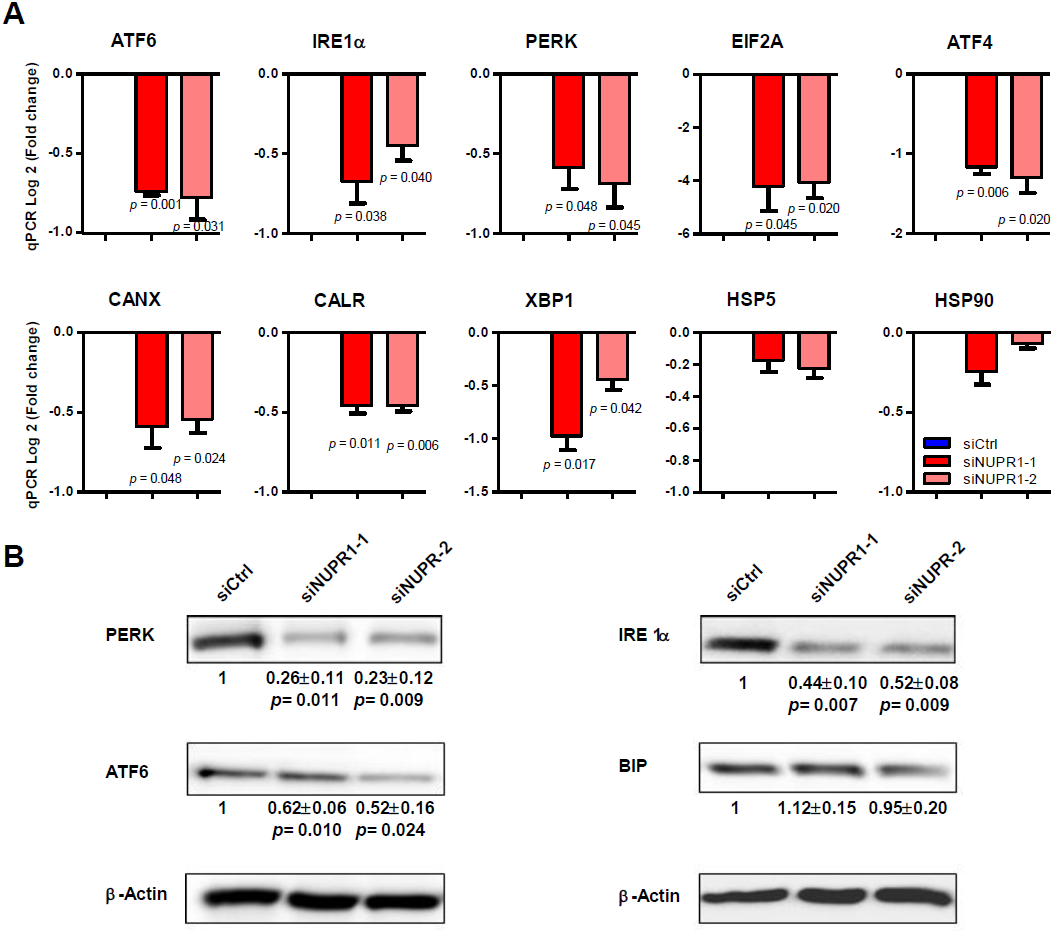
*NUPR1* deficiency induces a decrease in the expression of ER stress genes. Total RNA was extracted from MiaPaCa2 cells transfected with siCtrl, siNUPR1-1 or siNUPR1-2 for 72 h to evaluate the mRNA levels of the genes involved in ER stress response using RT-qPCR (A). Western blot analysis was performed to evaluated protein levels of expression of IRE1α, ATF6, PERK and BIP in NUPR1-depleted cells. Data are means of triplicates ± SEM. Statistically significant differences (p values ≤ 0.05) from siCtrl are shown.

### NUPR1 expression protect acinar cells of necrosis in mice with acute pancreatitis

During the course of pancreatitis, there is a sustained ER stress and pathological calcium signaling (Sah et al, 2014). Thus, we analyzed the effects of NUPR1 inactivation on cell death induced by experimental pancreatitis in *Nupr1-knockout* (*KO*) and *WT* mice. As shown in Figures 7A and B, we found that acute pancreatitis, induced by repeated injections of caerulein, resulted in a higher serum concentration of lipase (i.e.: 1248 ± 39 U/L and 715 ± 45 U/L after 12 h of induction in *KO* and *WT* mice respectively; p=0.0009) and lactate dehydrogenase (LDH) (i.e.: 4247 ± 198 ng/ml and 2034 ± 119 ng/ml after 12 h of induction in *KO* and *WT* mice respectively; p=0.002). Congruent with a role of NUPR1 in cell death by necrosis in acute pancreatitis, *Nupr1-KO* mice displayed a higher necrotic score when compared with wild type littermates (Figure 7C). Thus, this *in vivo* experiment demonstrates that normal levels of NUPR1 are necessary for protecting pancreatic cells from necrotic cell death.

**Figure 7.**
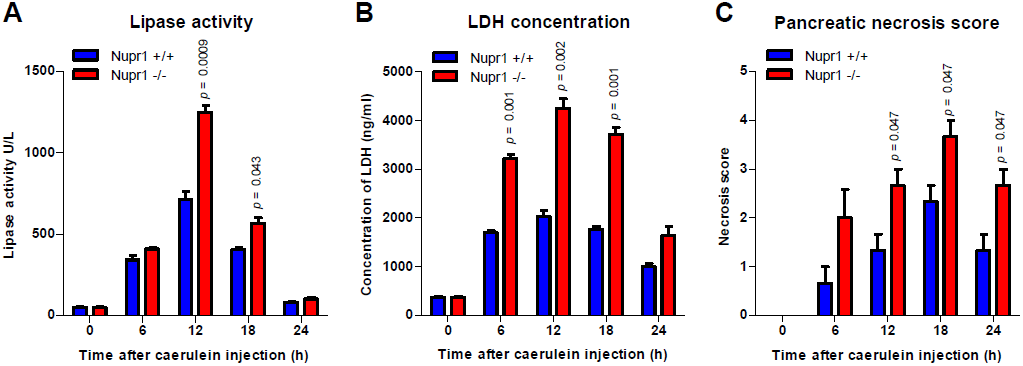
Nupr1 protects from pancreatitis-induced cell death. Serum lipase (A) and LDH (B) levels were measured in control Nupr1^+/+^ and Nupr1^−/−^ mice at 6, 12, 18, and 24 h following caerulein-induced acute pancreatitis. The necrosis score (C) was evaluated in histologic sections of pancreases and it refers to ranges of the percentages of cells involved: 0 = 0-5%; 1 = 5-15%; 2 = 15-35%; 3 = 35-50%; 4 ≥ 50%. Data are means of triplicates ± SEM. For each treatment, statistically significant differences (only p values ≤ 0.05) from Nupr1^+/+^ mice are shown.

## Discussion

The current study significantly contributes to a better understanding of mechanisms underlying the regulation of cell death by the cancer-associated protein, NUPR1. Our data support the idea that NUPR1 activation is strongly involved in protecting the cell against programmed cell death by both apoptosis and necrosis, and therefore, targeting NUPR1 protein in cancer appears to be a promising tool as an anticancer therapeutic strategy. Several empiric observations have shown that cancer cells undergo growth arrest or cell death after genetic inactivation of NUPR1 *in vitro* and *in vivo* (Emma et al, 2016; Guo et al, 2012; Kim et al, 2012; Li et al, 2017; Sandi et al, 2011; Vasseur et al, 2002b). However, to date, the mechanisms responsible for the biological consequences of its inactivation in cancer cells remain poorly understood. In this work, we used a battery of complementary methodological approaches to study these mechanisms, including *Nupr1-KO* mice, two siRNA sequences and CRISPR-Cas9 to inhibit NUPR1 expression with systematically concordant results. Notably, inactivation of this protein prevents pancreatic cells from properly responding to ER stress induced *in vitro* and leads to spontaneous metabolic cell death. We demonstrate that the inactivation of *Nupr1* in pancreatic cells promotes a cascade of events with several important consequences. For instance, loss of NUPR1 results in lower ATP production, higher consumption of glucose, a partial switch from OXPHOS to glycolysis, and a perinuclear relocalization of mitochondria to the vicinity of the ER. We also found that NUPR1 inactivation triggers a massive release of Ca^2+^ from the ER to the cytosol and a strong increase in ROS production, even though mitochondria produced less ATP. Altogether, these data enable us to describe a model in which inactivation of NUPR1 in pancreatic cancer cells results in an inefficient ER stress response through the inappropriate expression of several ER stress-associated genes. However, this adaptive attempt of the cell is counteracted by the increased Ca^2+^ concentration in the cytoplasm, which as shown previously (Berridge, 2002), leads to an increased mitochondrial Ca^2+^ concentration, disruption of mitochondrial membrane potential, leak of electrons from the respiratory chain in the complexes I and III (Brookes et al, 2004), ultimately resulting in a ROS accumulation, and, most importantly, in lower ATP generation. The low level of ATP production by NUPR1-deficient cells seems to be partially compensated by switching the remaining glucose consumption to glycolysis. However, this is insufficient to survive and therefore a metabolic cell death ensues.

NUPR1 was previously described as a protective factor against pancreatitis-induced lesions by activating expression of the anti-inflammatory factor pancreatitis associated protein I (also known as PAPI or Reg3beta) (Vasseur et al, 2004), in part through regulating its transcriptional activation. In turn, PAPI/Reg3beta inhibits the activation of NFkappaB in macrophages and therefore acts as a powerful anti-inflammatory factor (Folch-Puy et al, 2006). Consequently, the anti-inflammatory effect of NUPR1 could be explained through the activation of the PAPI/Reg3beta. Surprisingly, in this work we are demonstrating that experimental pancreatitis induced in NUPR1-deficient mice results in a significant increase in necrosis as indicated by the greater release of LDH, amylase and lipase in blood during the acute phase of the disease and by the significant increased necrotic score. Altogether, these results strongly suggest that induction of NUPR1 during acute pancreatitis controls both, inflammation and necrosis, by two different and complementary mechanisms.

A metabolism-associated role of NUPR1 was previously suggested in hepatocellular carcinoma, in which expression of *Nupr1* is induced in response to the mitochondrial malfunction, and consequently, NUPR1 overexpression favors cancer progression (Lee et al, 2015a; Lee et al, 2015b). In addition, it was also demonstrated that NUPR1 overexpression is necessary in the liver for an optimal response to a non-essential aminoacids-deficient diet and in pancreatic cancer cells to a starved culture media (Hamidi et al, 2012a; Maida et al, 2016). Based on these observations, we can assume that overexpression of NUPR1 operates to improve mitochondrial function to produce a maximum of energy in the presence of low substrates under stressed conditions.

Finally, we previously demonstrated that deficiency in NUPR1 expression results in induction of cellular cannibalism (Cano et al, 2012) and autophagy (Kong et al, 2010) processes. In light of the results from the current study, it is tempting to speculate that both cannibalism and autophagy could serve to generate energy fuel in NUPR1-deficient cells to compensate for the ATP deficiency. In addition, since intracellular ATP deficiency is accompanied by an increase in AMP and therefore activation of AMPK, it is possible that this kinase signals the initiation of the autophagy (Russell et al, 2014) and cellular cannibalism processes (Poels et al, 2012).

In conclusion, this current work demonstrates that, in NUPR1-deficient cells, glucose consumption is switched towards glycolysis instead of OXPHOS, which in turn results in a poor ATP production, and the natural promotion of a programmed necrotic process that we termed metabolic cell death. This defect is due to a mitochondrial malfunction, which in turn results as a consequence of a deficient ER stress response induced by the lack of NUPR1 expression.

### Materials and methods

#### Animals

*Nupr1*^−/−^ mice bear a homozygous deletion of exon 2 and have been reported previously (Vasseur et al, 2002a). The *Pdx1-cre; LSL-Kras^G^12^D^* mice were provided by R. Depinho (Dana-Faber Cancer Institute, Boston, Massachusetts, USA) and resulted from crossbreeding of the *Pdx1-Cre* and *LSL-Kras^G12D^* strains. Mice were kept in the Experimental Animal House of the Centre de Cancérologie de Marseille (CRCM) of Luminy. All experimental procedures on animals were approved by the Comité d’é thique de Marseille numé ro 14 and in accordance with the European Union regulations for animal experiments. **Induction of experimental pancreatitis**: Pancreatitis was induced in 4-month-old *Nupr1*^−/−^ and *Nupr1*^+/+^ mice weighing 20-24 g. Caerulein (Sigma) was administered as intraperitoneal injections of 50 μg/kg body weight, as previously described (Halangk et al, 2000). Saline-injected animals served as controls.

#### Preparation of isolated pancreatic acinar cells

Briefly, 5-week-old mice pancreata were immediately removed after euthanasia, cut into small sections, and digested with collagenase IA (Sigma-Aldrich) for 10 min at 37°C as previously described (Santofimia-Castano et al, 2015). Acini were collected by centrifugation at 30 g for 5 min, washed and resuspended in Seahorse medium, and plated in pre-coated Cell Tak (Corning) Seahorse plates (Agilent).

#### Preparation of serum and tissue samples

Mice were sacrificed at several time points from 3 to 24 h following the first intraperitoneal injection of cerulein. Whole blood samples were centrifuged at 4°C was removed on ice, immediately frozen in liquid nitrogen, and stored at −80°C.

#### Necrosis score evaluation

Formalin-fixed samples were embedded in paraffin and 5-μm sections were stained with hematoxilin and eosin. The histological alterations in the sections were scored by three experienced reviewers. The scores refer to ranges of percentages of cells involved from 0 (absent), 1 (minimal, 0 to 15%), 2 (moderate, 15 to 35%), 3 (important, 35 to 50%) to 4 (maximal, >50%).

#### LDH and lipase serum concentration

A Cobas analyzer was used to measure enzyme concentrations in the serum samples of *Nupr1*^−/−^ and *Nupr1*^+/+^ mice, according to the manufacturer’s specifications and using proprietary reagents.

#### RNA-seq

RNA from pancreata of 5-week-old mice was purified following the procedure of Chirgwin et al. (Chirgwin et al, 1979). RNA integrity was assessed using the Agilent 2100 Bioanalyzer with the RNA 6000 Nano chip Kit. Samples were sent to the Transcriptomic and Genomic platform of Marseille-Luminy (TGML). The enrichment of mRNA, cDNA library construction for RNA-seq, sequencing, and bioinformatic data analysis were performed as previously described (Amrani et al, 2014).

#### Cell culture

MiaPaCa2 and Panc1 cells were obtained from the American Type Culture Collection and maintained in Dulbecco’s modified Eagle’s medium (DMEM) (Gibco, Life Technologies) supplemented with 10% fetal bovine serum (Lonza) at 37°C with 5% CO_2_.

#### siRNA transfection

Cells were plated at 70% confluence and INTERFERin reagent (Polyplus-transfection) was used to perform siRNA transfections, according to the manufacturer’s protocol. Scrambled siRNA that targets no known gene sequence was used as a negative control. All assays were carried out 72 h post-transfection. The sequences of Nupr1-specific siRNAweresiNUPR1-1r(GGAGGACCCAGGACAGGAU)dTdT and siNUPR1-2 r(AGGUCGCACCAAGAGAGAA)dTdT.

#### CRISPR-Cas9 clones development

Cells were seeded in 12-well plates and transfected with 1 μg of *Nupr1* double nickase plasmid or double control nickase plasmid (Santa Cruz Biotechnology), using Lipofectamine 3000 Transfection Reagent (Thermo Fisher Scientific) in each well, following the manufacturer’s protocol. Transfected cell selection was performed with puromycin and, after 72 h, single cell colonies were isolated. Complete allelic knockouts were confirmed using Sanger sequencing.

#### NUPR1 expression vector transfection

Panc1 cells (clones C1, NUPR1-KO1 and NUPR1-KO2) were seeded in 12-well plates and transfected with 2 μg of DNA (NUPR1-FLAG or control vector) and 2 mL of Lipofectamine 3000 Transfection Reagent (Thermo Fisher Scientific) per well. Cells were assayed after 24 h post-transfection.

#### LDH assay, ATP production, and caspase-3/7 activity assay

Cells were seeded at a density of 10,000 cells per well in 96-well plates. Cells were allowed to attach overnight and treated the next day for 24 h. At the end of the experiment, LDH release, ATP production, and caspase-3/7 activity were monitored using CytoTox-ONE (Promega G7890), CellTiter-Glo (Promega G7571) and Caspase-Glo 3/7assay (Promega G8091), respectively. Data were normalized to the cell number.

#### RT-qPCR

RNA from cells was isolated using Trizol reagent (Invitrogen) and reverse-transcribed using Go Script (Promega) according to the manufacturer’s instruction. Real-time quantitative PCR (RT-qPCR) was performed in a Stratagene MXPro-MX3005P using Promega reagents. Primer sequences are available upon request.

#### Western blot

Proteins were resolved by SDS-PAGE, transferred to nitrocellulose filters, revealed with ECL and detected using a Fusion FX7 imager (Vilber-Lourmat, France). The following antibodies were used: rabbit monoclonal anti-PERK (C33E10, CTS), rabbit polyclonal anti-BIP (C50B12, CTS), rabbit polyclonal anti-IRE1α (14C10, CTS), rabbit monoclonal anti-ATF6 (D4Z8V, CST), rabbit polyclonal anti-human NUPR1 antibody (home made) and anti-β-actin (A5316, Sigma).

#### Extracellular flux assay

Measurements were performed using a Seahorse Bioscience XF24 Extracellular Flux Analyzer. This system allowed measurement of the cellular oxygen consumption rate (OCR, in pmoles/min), the extracellular acidification rate (ECAR in mpH/min) and the proton production rate (PPR). Cells were plated at 30,000 cells/well onto Seahorse 24-well plates 48 h before the assay. OCR was measured using the XF Cell Mito Stress Test Kit under basal conditions, and in response to 1μM oligomycin and 0.25 or 0.5μM of carbonylcyanide p-(trifluoro-methoxy) phenylhydrazone (FCCP) in MiaPaCa2 or pancreatic acini, respectively. Oligomycin injection allowed calculation of the OCR for ATP production, and FCCP treatment gave two indexes; the maximal OCR capacity and the spare respiratory capacity. Finally, OCR was stopped by adding the electron transport chain inhibitors rotenone and antimycin A (0.5μM each). ECAR was measured using the Seahorse XF Glycolysis Stress Test Kit. Glycolysis was calculated as the difference between the values of ECAR upon glucose addition (10 mM) and basal ECAR. Glycolytic capacity was calculated as the difference between ECAR following the injection of 1 μm oligomycin, and the basal ECAR. Finally, glycolytic reserve was calculated as the difference in ECAR between glucose and oligomycin injections. At the end of the experiment, glycolysis was stopped by adding 2-Deoxyglucose (100 mM). The contribution of OXPHOS and glycolysis to ATP production was calculated using the OCR and PPR as previously described (Wu et al, 2016). The Seahorse XF Mito Fuel Flex Test was used to determine the rate of oxidation of each fuel by measuring OCR. Inhibitors of glutaminase (BPTES 3 μM), carnitine palmitoyl-transferase 1A (Etomoxir 4 μM), and glucose oxidation (UK5099 2 μM) were used. Levels of OCR and ECAR were normalized to the cell number.

#### Lactate release, and glucose and glutamine uptake

The culture media was examined for levels of glucose, glutamine, and lactic acid using the YSI 2900 Biochemistry Analyzer (YSI Life Sciences, Yellow Springs, OH, USA) following the manufacturer’s instructions. The increase in lactate production was calculated by subtracting the lactate present in the control medium (without exposure to cells) from the lactate present in the experimental media. Glucose and glutamine uptake were calculated by subtracting the level of the experimental media from the level of the control medium.

#### AnnexinV/PI staining

Cells were collected after incubation for 24 h with the stimuli. Cells were washed with PBS, then detached with Accutase (Gibco, Life Technologies), and resuspended in 100 μl of 1X Annexin-binding buffer. Pacific-Blue Annexin V (5 μl, BioLegend) was added to the cell suspension and after a 15-min incubation at room temperature, 400 μl of 1X Annexin-binding buffer was added. Before analysis by flow cytometry, propidium iodide (5 μl, Miltenyi Biotec) was added to the suspension. 10,000 events per sample were collected in a MACSQuant-VYB (Miltenyi Biotec, Surrey, UK). Data analysis was performed using the FlowJo software.

#### Superoxide anion production by the mitochondria

MitoSOX (Molecular Probes) was added to a final concentration of 5 μM for 10 min. After incubation, cells were washed with warm PBS, detached with Accutase, and resuspended in HBSS (Gibco, Life Technologies) for flow cytometry; or fixed with 4% paraformaldehyde for 10 min at room temperature and mounted using Prolong Gold antifade reagent with DAPI (Invitrogen). In cytometry experiments, 10,000 events per sample were collected in a MACSQuant-VYB and data analysis was performed using FlowJo software. Image acquisition of mounted samples was carried out on a microscope Nikon Eclipse 90i fluorescence microscope.

#### Mitochondrial membrane potential

Measurement was performed using the MITO-ID Membrane potential detection kit (ENZ-51018) according to the manufacturer’s protocol. Briefly, cells were collected, washed, and preincubated in 500 μl of the Assay Solution containing 5 μl of MITO-ID MP Detection Reagent for 15 min. After this, 10,000 events per sample were collected in a MACSQuant-VYB and orange fluorescence data analysis was performed using the FlowJo software.

#### Cytosolic calcium concentration

Cells were preincubated with Fluo-4 AM (Molecular Probes) for 30 min at 37°C. Then, cells were washed and resuspended in HBSS for flow cytometry analysis. 10,000 events per sample were collected in a MACSQuant-VYB and data analysis from flow cytometry was performed using FlowJo software.

#### Mitochondrial network

Localization of the mitochondrial network was assayed by incubation of cells in the presence of MitoTracker DeepRed FM (200 nM, Molecular Probes) at 37°C for 30 min. Then, cells were washed twice with PBS and fixed with 4% paraformaldehyde. Cells were subsequently permeabilized, incubated with 1% BSA and after, with Alexa Fluor 488 Phalloidin, according to the manufacturer’s instructions. Finally, samples were mounted using the Prolong Gold antifade reagent with DAPI. Confocal images were acquired using an inverted microscope equipped with LSM 880 with Airyscan detector controlled by Zeiss Zen Black, 63x lens.

#### Electron microscopy

Samples were prepared using the NCMIR protocol for SBF-SEM (Deerinck et al, 2010). 70 nm-ultrathin sections were prepared using a Leica UCT Ultramicrotome (Leica, Austria) and deposited on formvar-coated slot grids. The grids were observed in an FEI Tecnai G2 at 200 KeV and acquisition was performed on a Veleta camera (Olympus, Japan).

#### DNA microarray

Total RNA (15 μg) was isolated and reverse transcribed for hybridization to the human oligonuleotide array U133 Plus 2.0 (Genechip, Affymetrix) as described previously (Hamidi et al, 2012b). Arrays were processed using the Affymetrix GeneChip Fluidic Station 450 and scanned using a GeneChip Scanner 3000 G7 (Affymetrix). The GeneChip Operating Software (Affymetrix GCOS v1.4) was used to obtain chip images with quality control conducted using the AffyQCReport software.

#### Statistics

Statistical analyses were performed using the unpaired 2-tailed Student t test and non-normal distribution. A p value less than or equal to 0.05 was considered significant. All values are expressed as mean ± SEM. RT-qPCR data are representative of at least three independent experiments with technical triplicates completed.

## Acknowledgments

This work was supported by La Ligue Contre le Cancer, INCa, Canceropole PACA, DGOS (labellisation SIRIC) and INSERM. PS-C is supported by Fundación Alfonso Martí n Escudero and Fondation de France. The electron microscopy experiments were performed in the PiCSL-FBI core facilty (IBDM, AMU-Marseille). The confocal microscopy experiments were performed in the “Plateforme de Microscopie et Imagerie Scientifique” of CRCM of Marseille.

## Author contributions

PS-C, PS, and JLI proposed the concept and designed the experiments; PS-C, WL, JB, OG, and GL carried out experiments; PS-C, GL, AC, JLN, AG, RU, PS, and JLI analyzed and interpreted the results; PS-C, GL, JLN, AG, RU, PS, and JLI wrote the manuscript.

## Legend of supplementary Figures

**Figure S1.**
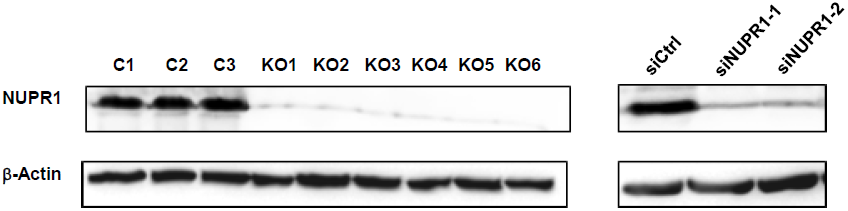
NUPR1 protein expression. Western blot analysis was performed in 3 control and 6 CRISPR-Cas9 NUPR1 knockout Panc-1 clones (right panel) and in MiaPaCa2 cells transfected with siCtrl and siNUPR1-1 and siNUPR1-2 (left panel) to evaluate the NUPR1 protein levels.

**Figure S2.**
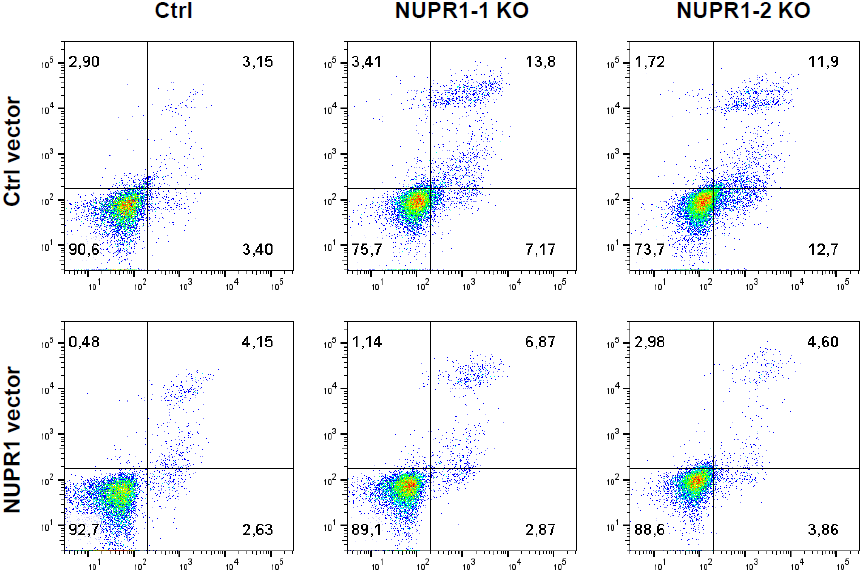
Reconstitution of NUPR1 expression in CRISPR-Cas9 knockout Panc-1 cells reduces cell death. Flow cytometry analysis was carried out after Annexin-V and PI staining in 2 independent Panc-1 *Nupr1* knockout clones, transfected with NUPR1-FLAG or a control vector. A representative experiment of the dot plot profile of cells was showed.

**Figure S3.**
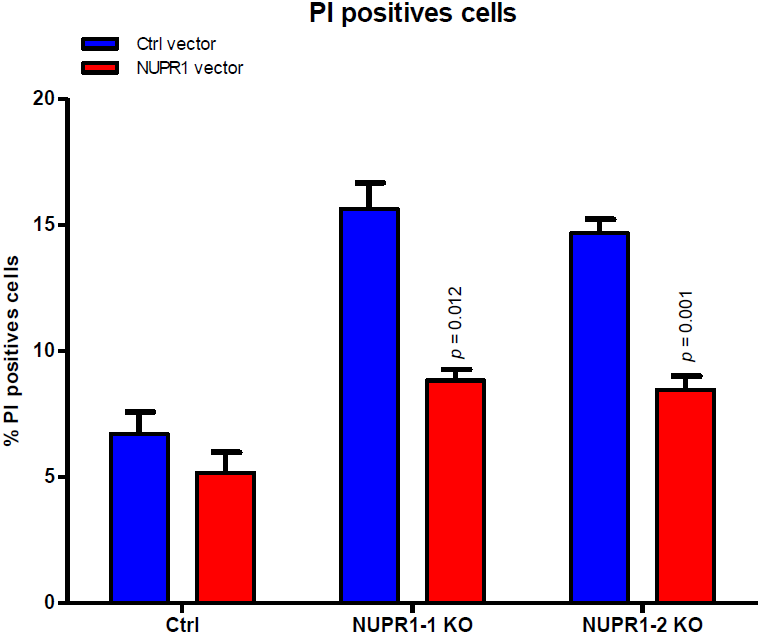
Reconstitution of NUPR1 expression in CRISPR-Cas9 knockout Panc-1 cells reduces cell death. Flow cytometry analysis was carried out after Annexin-V and PI staining in 2 independent Panc-1 *Nupr1* knockout clones, transfected with NUPR1-FLAG or a control vector. Data are means of triplicates ± SEM of 1 control and 2 *Nupr1* knockout clones. Statistically significant differences (p values ≤ 0.05) are shown.

